# Phenology across scales: an intercontinental analysis of leaf-out dates in temperate deciduous tree communities

**DOI:** 10.1101/2023.11.21.568089

**Authors:** Nicolas Delpierre, Suzon Garnier, Hugo Treuil-Dussouet, Koen Hufkens, Jianhong Lin, Colin Beier, Michael Bell, Daniel Berveiller, Matthias Cuntz, Giulio Curioni, Kyla Dahlin, Sander O. Denham, Ankur R. Desai, Jean-Christophe Domec, Kris M. Hart, Andreas Ibrom, Emilie Joetzjer, John King, Anne Klosterhalfen, Franziska Koebsch, Peter Mc Hale, Alexandre Morfin, J. William Munger, Asko Noormets, Kim Pilegaard, Felix Pohl, Corinna Rebmann, Andrew D. Richardson, David Rothstein, Mark D. Schwartz, Matthew Wilkinson, Kamel Soudani

## Abstract

**Aim:** To quantify the intra-community variability of leaf-out (ICVLo) among dominant trees in temperate deciduous forests, assess its links with specific and phylogenetic diversity, identify its environmental drivers, and deduce its ecological consequences with regard to radiation received and exposure to late frost.

**Location:** Eastern North America (ENA) and Europe (EUR).

**Time period:** 2009-2022

**Major taxa studied:** Temperate deciduous forest trees.

**Methods:** We developed an approach to quantify ICVLo through the analysis of RGB images taken from phenological cameras. We related ICVLo to species richness, phylogenetic diversity and environmental conditions. We quantified the intra-community variability of the amount of radiation received and of exposure to late frost.

**Results:** Leaf-out occurred over a longer time interval in ENA than in EUR. The sensitivity of leaf-out to temperature was identical in both regions (-3.4 days per °C). The distributions of ICVLo were similar in EUR and ENA forests, despite the latter being more species-rich and phylogenetically diverse. In both regions, cooler conditions and an earlier occurrence of leaf-out resulted in higher ICVLo. ICVLo resulted in a ca. 8% difference of radiation absorption over spring among individual trees. Forest communities in ENA had shorter safety margins as regards the exposure to late frosts, and were actually more frequently exposed to late frosts.

**Main conclusions:** We conducted the first intercontinental analysis of the variability of leaf-out at the scale of tree communities. North American and European forests showed similar ICVLo, in spite of their differences in terms of species richness and phylogenetic diversity, highlighting the relevance of environmental controls on ICVLo. We quantified two ecological implications of ICVLo (difference in terms of radiation absorption and exposure to late frost), which should be explored in the context of ongoing climate change, which affects trees differently according to their phenological niche.

## 1 Introduction

### 1.1 Intra-community variability of leaf phenology in temperate forests

The phenology of the tree canopy strongly influences the functioning of forests (Barr et al., 2007; Delpierre et al., 2009; Richardson et al., 2010) and of the climate system (Richardson et al., 2013) by modulating the exchange of matter and energy with the atmosphere. A wealth of study has been devoted to identify the environmental and biological drivers of spring leaf-out. These studies have highlighted the central roles of temperature and photoperiod (see Delpierre et al., 2016 for a review). Almost all these studies have focused on the average date of leaf-out in the ecosystem. Yet, when conducting phenological observations in a forest, one can observe a large inter-individual variability of leaf-out among conspecifics. In a preceding study, we showed that the average variability of leaf-out within a population of trees is 19 days, which corresponds to 75% of the variability observed at the scale of the continent (considered species were temperate oaks and beech, see Delpierre et al., 2017).

Such a wide range in leaf-out date could arise from phenology being a neutral trait for the tree, not affecting its fitness and therefore not being the object of natural selection. This idea is currently an object of debate, with little data documenting the link between phenology and fruit productivity (as a direct measure of fitness). Some studies have investigated the link between phenology and growth (an indirect measure of fitness) but their results are partly contradictory, with some showing no link between interannual variations of ring width indices and leaf phenology for a given tree (Cufar et al., 2015; de Sauvage et al., 2022), and others showing a globally significant, but not systematic, link between basal area increment and leaf phenology among dominant conspecifics in a given population (Delpierre et al., 2017). Leaving that debate aside, it is likely that the wide range of leaf-out dates observed in forests is due to the process of stabilising selection in which environmental conditions will impose limits to the acceptable variability achievable in phenological traits, while within-community interactions will favor inter-individual variability (Violle et al., 2012). In that context, fluctuating interactions and hazards may favor a large variability of phenological traits in a population (Alberto et al., 2011). For instance, individual trees that leaf-out late will probably be advantaged in years with a late frost, but may logically be disadvantaged in years when early spring conditions are favorable, or when pathogens flourish (Dantec et al., 2015). Ongoing climate change is accompanied by changes in the probability of exposure to late frost (Zohner et al., 2020), which could influence communities differently in areas where the probability is increasing (e.g. Europe, but see Lin et al., prep) than in areas where it is decreasing (e.g. North America).

The factors determining the intra-population variability of leaf-out have been little studied, but it is established that edaphic conditions (nature of the soil in Arend et al., 2016, soil water content in Delpierre et al., 2017), microclimate (e.g. local seasonal air temperature in Donnelly et al., 2017), genetic variability (Bontemps et al., 2015) and ontogeny (Vitasse, 2013) are involved. Furthermore, the intra-population variability in leaf-out itself varies between years, depending on the prevailing micrometeorological conditions. Thus, intra-population variability is all the more marked the colder the temperature conditions during leaf-out (Deńechère et al., 2021; Delpierre et al., 2020).

The amplitude of leaf-out (i.e. the duration from the earliest to latest tree to leaf out) is likely to increase as one moves from the population to the community, encompassing a wider range of phenological niches. These niches are distributed vertically (e.g. dominant vs. understory tree species, Richardson and O’Keefe, 2009) and horizontally (e.g. early vs. late species in the overstory, Cole and Sheldon, 2017). The evolutionary history of species (i.e. genetic determinism) explains the inter-specific differences in leaf-out within a community. In this context, a more species-rich community would also be expected to have a greater phenological range, the phenological range of the community being composed of the specific phenological ranges (Fig. S1). In addition to species richness, the phylogenetic diversity of communities deserves to be considered, as plants display a certain phylogenetic conservatism (whereby phylogenetically close species display similar phenological traits, Davies et al., 2013; Panchen et al., 2014). For example, one would expect a larger intra-community variability of leaf-out (ICVLo) in North American forests than in European forests, all else being equal, because they are more species-rich (Liang et al., 2022; Latham and Ricklefs, 1993) and display a higher phylogenetic diversity (Eiserhardt et al., 2015). The ICVLo is itself susceptible to variations from year to year (Fig. S1). By extending the results obtained at the population level (Deńechère et al., 2021) to the community, we hypothesize that cold temperatures during the leaf-out period would increase the ICVLo.

### 1.2 Using phenocams to study the intra-community variability of leaf phenology

A large part of the literature on forest leaf phenology is based on ground observation data. These data are historically the oldest and have been collected in national to continental databases (e.g. PEP725 Templ et al., 2018, NPN-usa Betancourt et al., 2007) that cover a period of several decades (e.g. from the 1950’s in the PEP725 database). In the late 2000’s, the use of ”digital repeat photography” to document the phenology of plant canopies became widespread (Richardson et al., 2007, 2009). These studies were initially based on the analysis of data obtained by automated photographic instruments, most often mounted on towers overhanging the canopy (i.e. phenological cameras or ”phenocams”). Networks of phenocams have been set up (notably the PhenoCam Network, Richardson, 2019, see also Wingate et al., 2015), which has enabled data to be centralized and harmonized. Image data are more complex than ground observation data.

They have to be processed to extract an analysable phenological signal (e.g. critical dates in the development of the foliage). The development and public sharing of algorithms for processing phenocam images (e.g. R packages *phenopix* Filippa et al., 2016 and *xROI* Seyednasrollah et al., 2019b) has increased the use and impact of these data. Several studies have been dedicated to the comparison of critical phenological dates observed from the ground and inferred from the analysis of phenocam datasets at a common site (e.g. Keenan et al., 2014; Soudani et al., 2021). They show a very good match between ground-observed and phenocam-inferred leaf-out dates at the community scale. More recently, we have shown by comparing ground observation and phenocam data that the analysis of phenocam data also allows quantifying the intra-population variability of leaf-out (Delpierre et al., 2020). For this purpose, we subdivided the phenocam scene (i.e. Region Of Interest, ROI at the canopy scale) into several sub-ROIs (each targeting a particular tree) and analysed the data at this scale. The idea of analysing intra-canopy (i.e. intra-community) variability in bud break is not new, and had previously appeared in site-scale studies (Ahrends et al., 2008; Richardson et al., 2009; Filippa et al., 2016). To our knowledge, it has not been deployed in the context of a regional study yet.

### 1.3 Objectives

Here, we analysed phenocam data obtained over North American and European forests with the aim to investigate the determinants and ecological consequences of ICVLo in temperate deciduous forests. Specifically, we answered the following four questions:

1. Do variations in spring temperature influence similarly the date of leaf-out in North American and European temperate deciduous forests?
2. Is the higher species richness and/or phylogenetic diversity of North American forests associated with a higher ICVLo?
3. What are the environmental controls of ICVLo?
4. What are the ecological implications of ICVLo, in terms of light absorption and exposure to late frosts?

## 2 Materials and Methods

### 2.1 Study sites

We analysed images taken by phenological cameras over 17 sites located in Europe (EUR, 8 sites) and Eastern North America (ENA, 9 sites) (Fig. 1, Table 1). The sites were distributed between 36°N to 56°N, and classified into Köppen-Geiger climate zones using the *kgc* package (Bryant et al., 2017), with zone names from Kottek et al., 2006. We selected these sites on the basis of the length of the image time series available and the general quality of the images (main criteria were : fixed field of view including a large proportion of forest cover, and good image sharpness). The quality of the images was not always consistent between years for the same site, which led us to keep some years and discard others for a given site (Table 1). Analysed images were acquired from 2009 to 2022.

**Fig. 1:**
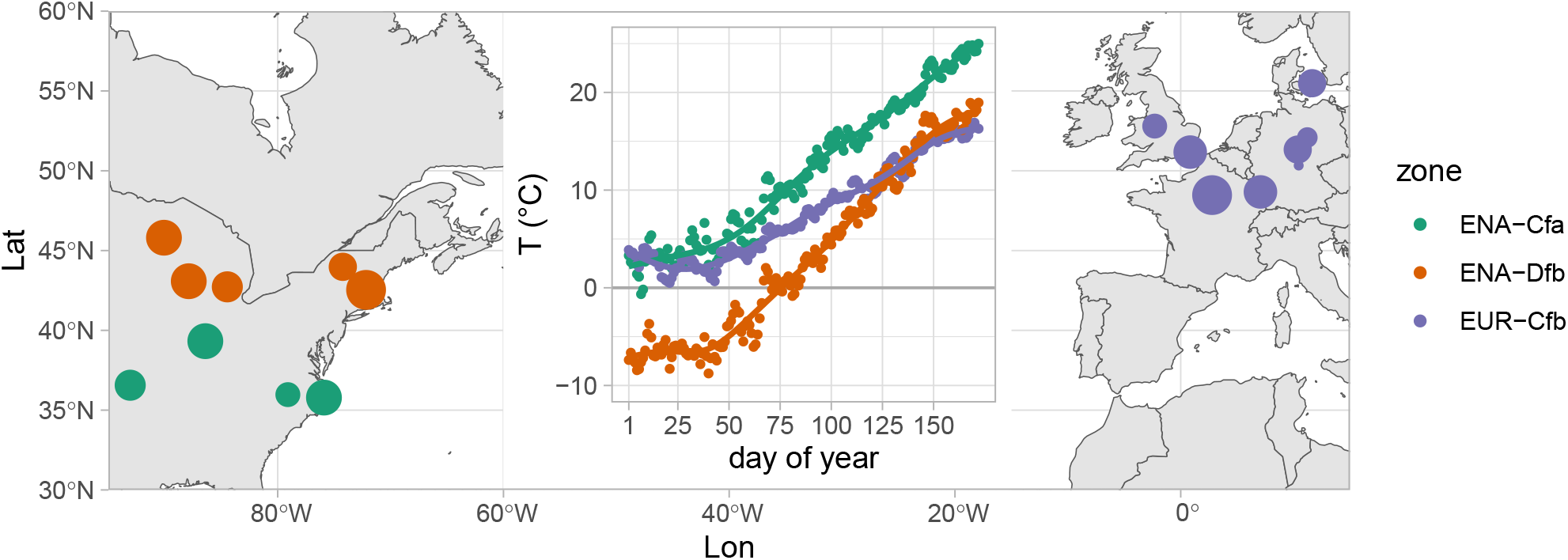
Location of study sites. Colors indicate classification into Köppen climate zones. The size of points on the map scales with the number of years, ranging from 2 (smallest) to 10 (largest). The inset graph represents daily average air temperature climatologies (T, in °C) established over January to June; temperature data were binned according to day of year (DoY) and averaged across all study site-years of the corresponding climate zone. Climate zones are: Cfa = warm temperate with hot summer, Cfb = warm temperate with warm summer, Dfb = snow with warm summer.

**Table 1:**
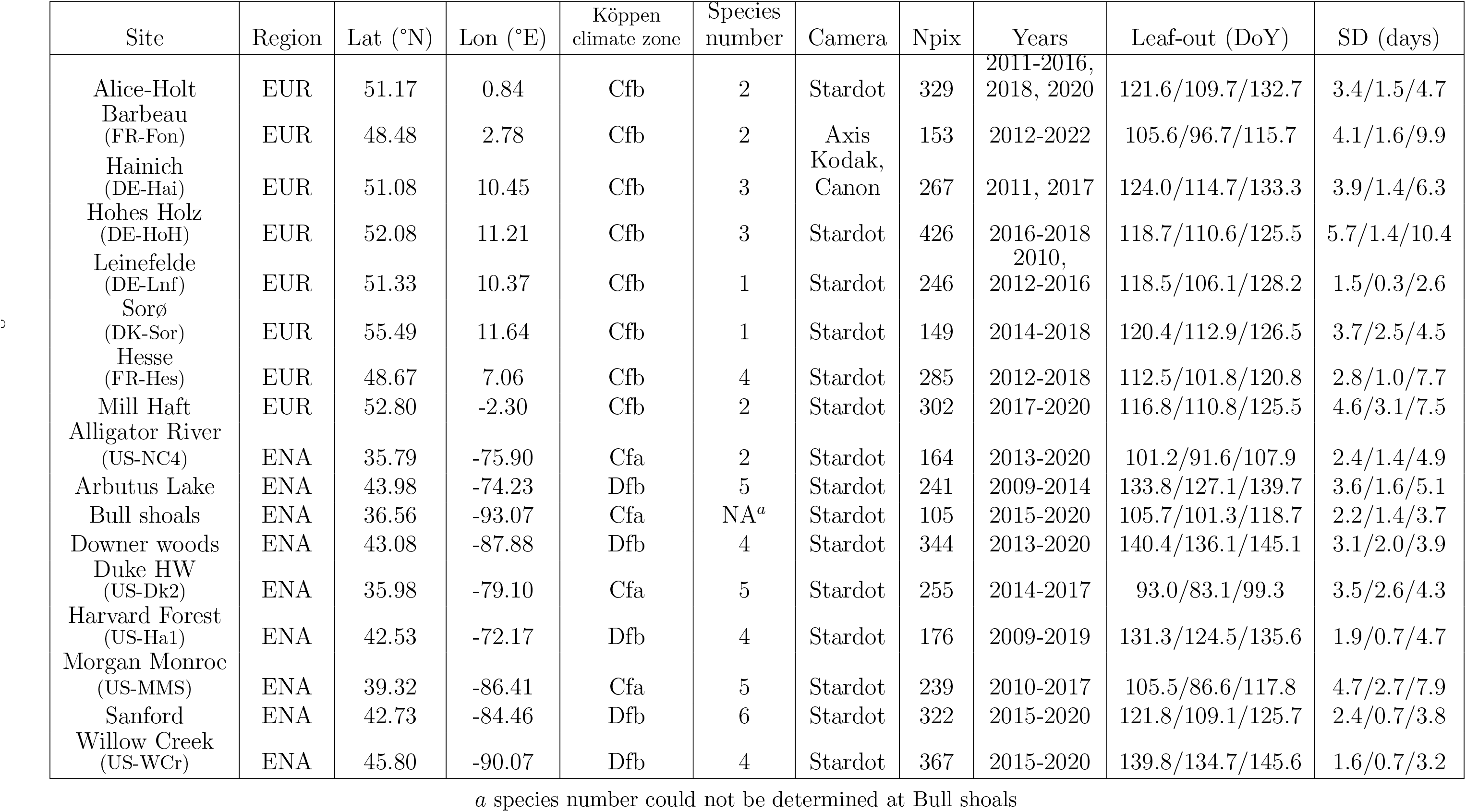
Characteristics of study sites. *Npix* refers to the number of sub-ROIs analysed at each site. *SD* is the standard deviation of leaf-out dates. Columns *Leaf-out* and *SD* contain the mean / minimum / maximum values observed at each site across the measuring time period (Years). Köppen climate zones: Cfa = warm temperate with hot summer, Cfb = warm temperate with warm summer, Dfb = snow with warm summer. *Stardot* is for Stardot NetCam SC 5MP, *Axis* is for Axis P1347, *Kodak* is for Kodak DC290, *Canon* is for Canon Powershot A700. The actual species names are displayed in Table S1.

### 2.2 Processing phenocam images

We obtained images from the phenocam dataset (Seyednasrollah et al., 2019b) for 10 study sites (all the ENA sites plus Mill Haft). Images for the 7 other sites were provided by the site PIs. For each site, we delineated a mask (Region Of Interest, ROI) to delimit the deciduous vegetation zone in the phenocam scene (i.e. excluding roads, buildings, the sky and evergreen trees), thanks to the R package *xROI* (Seyednasrollah et al., 2019a). Since our aim was to work on ICVLo, we adopted a ”pixel-based” approach (Filippa et al., 2016), that has rarely been used to date for the analysis of phenocam images (but see Ahrends et al., 2008; Delpierre et al., 2020; Richardson et al., 2009). For this, we subdivided the ROI into sub-ROIs using a systematic hexagonal grid (Fig. 2) with the R package *rgeos* (Bivand and Rundel, 2023). The phenological sequence of each sub-ROI was analysed independently.

**Fig. 2:**
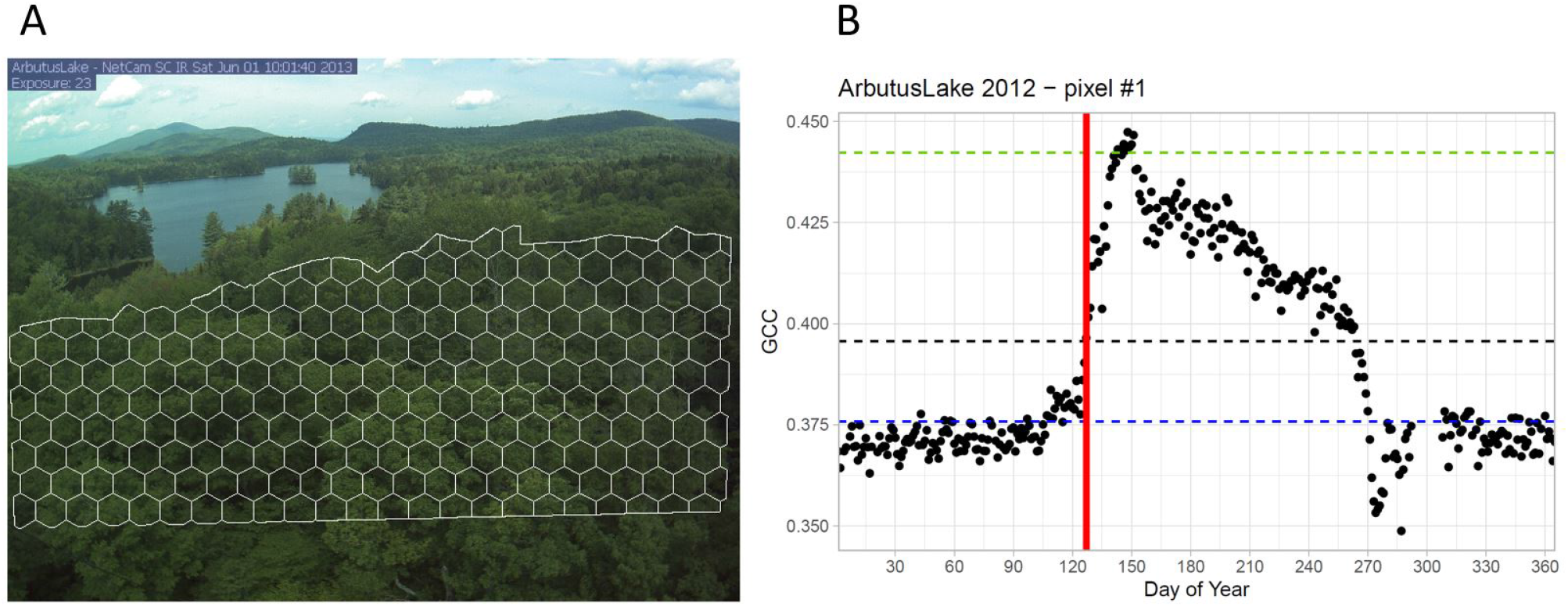
Processing of the phenocam data. (A) representation of a scene onto which a gridded Region Of Interest (ROI) has been applied; (B) data extracted from one of the grid elements (i.e. one sub-ROI), where the blue horizontal line marks the minimum spring Green Chromatic Coordinate (GCC) value, green horizontal line is the maximum GCC value considered, and the black horizontal line represents 30% of the amplitude between the blue and green lines. Red vertical line is the leaf-out date determined for this particular grid element. Images and data are from Arbutus Lake site (NY, USA) in 2012.

Using the sub-ROI approach in a previous work, we showed it was possible to retrieve intra-scene variability of leaf-out dates that were very close to the inter-tree variability of leaf-out observed from the ground (Delpierre et al., 2020). In our systematic grid approach, the mesh size was specifically chosen for each site according to the proximity and size of the tree crowns (Fig. 2A). We chose mesh sizes that were slightly smaller than the average tree crown size observed on the grid, approximating one sub-ROI to represent one tree. The idea here was to reduce the risk of an under-estimation of the intra-community variability of leaf-out that would result from choosing too coarse mesh sizes (i.e. that would lead to confound canopy crowns), bearing in mind that the intra-crown variability of leaf-out is lower than the inter-crown variability (e.g. Smith, 2018, their table A3.3). We conducted preliminary evaluation of the method on a subset of four sites, comparing the intra-scene variation of leaf-out obtained from the systematic grid vs. a more detailed approach into which we identified sub-ROI at the scale of the tree (Suppl. Notes S2). The error arising from the use of the systematic grid was of 0.36 days (see Fig. S2.2), which yields a signal-to-noise ratio of 8.6 (calculated as the ratio of the average standard deviation (SD) of leaf-out, see below, to the error of 0.36 days). We considered this value high enough to be confident in the quality of the analyses conducted on data obtained from the systematic grid approach.

### 2.3 Retrieving leaf-out from the GCC signal

In each sub-ROI, we determined the date of leaf-out from the analysis of the Green Chromatic Coordinate (GCC) time series (Keenan et al., 2014). GCC uses red (R), green (G), and blue (B) digital numbers to calculate the ratio of green within the image (GCC = G/(R+G+B)). Specifically, we determined the ”date of leaf-out” of each sub-ROI with a threshold approach (Keenan et al., 2014), using 30% of the spring signal amplitude as a threshold (Fig. 2B). We computed the lower bound of the signal amplitude as the 95th percentile of the GCC data obtained from day of year (DoY) 1 to 80 (blue line in Fig. 2B). We computed the upper bound of the signal amplitude as the 98th percentile of the GCC data obtained over the whole year (green line in Fig. 2B). After establishing the date of leaf-out for each sub-ROI, we first cross-checked the minimum and maximum dates of leaf-out against phenocam images. In eight site-years out of 106 site-years present in the dataset, the analysis of the GCC data produced too early or too late leaf-out dates in some sub-ROIs as compared to our visual inspections of the phenocam images. We removed those outliers from the distributions of leaf-out dates determined at the sub-ROI scale. These outliers represented 3% of the sub-ROIs on average for the eight site-years considered. Then, we computed community-scale phenological metrics for each site-year in the dataset, namely the minimum, maximum and average leaf-out date (DoY), the leaf-out standard deviation (in days) and the amplitude (i.e. maximum-minimum, in days). In the following, we use the standard deviation of leaf-out (SD, in days) for quantifying ICVLo. Standard deviation is a measure of the average duration between each sub-ROI leaf-out date and the average date established over the whole community.

### 2.4 Meteorological data

We retrieved air temperature and radiation data of each site from nearby meteorological stations for ENA sites (except Bull shoals, for which we used data from the DAYMET database (Thornton et al., 2021)) as well as for two European sites (Alice-Holt and Mill Haft), and from the ICOS community for the other European sites (”Warm Winter 2020 ecosystem eddy covariance flux product”, https://doi.org/10.18160/2G60-ZHAK). In order to assess the influence of temperature conditions on ICVLo, we computed for each site-year the average temperature (Tmean, in °C) and the absolute minimum temperature occurring during the period of leaf-out. Preliminary analyses (Fig. S3) identified that ICVLo was more strongly related to the minimum temperature measured during the period extending from the 5th to the 95th percentile of the sub-ROI distribution of leaf-out (hereafter Tmin, in °C).

In order to quantify how treewise light absorption was influenced by ICVLo, we compared the sum of radiation received by the community from (i) the leaf-out date of the earliest sub-ROI and (ii) the leaf-out date of the latest sub-ROI and October 1.

In order to compare the exposure of the earliest and latest trees in the community to late frosts, we quantified their respective ”safety margins”. For this purpose, we calculate the duration (in days) between the leaf-out date of the earliest and latest sub-ROI and the occurrence of minimum temperatures below the critical threshold of -3°C, below which frost damage on emerging leaves is irreversible (Lin et al., 2023).

### 2.5 Community diversity metrics

One of our objectives was to evaluate the influence of the community diversity on ICVLo, with the hypothesis that more diverse canopies would display higher ICVLo. For this, we considered two metrics for the diversity of the community. First, we considered the number of dominant tree species (*Spnum*) that were visible in the site ROIs. Yet, the species number could be less informative than phylogenetically-informed metric of the diversity, because leaf-out is a trait that is conserved in some clades (Panchen et al., 2014). Hence we also considered the *mean pairwise distance (MPD)* as a metric for community diversity. MPD is the mean phylogenetic distance (i.e. branch length) among all pairs of species within a community. Because all species composing a community do not have the same abundance in the analysed ROIs, we weighted the contribution of each species pair in MPD by the product of the individual species abundances (see Table S1). Calculations of MPD were done with the *picante* package (Kembel et al., 2010), using the tree phylogeny from Zanne et al., 2014. Since we could not identify the species identity and proportional contribution to ROI at Bull shoals, we included data from this site only in analyses that do not require these metrics (i.e. Fig. 3, Fig. 4A,C,D and Fig. 5 in the main text, Fig. S4 in the supplementary materials).

**Fig. 3:**
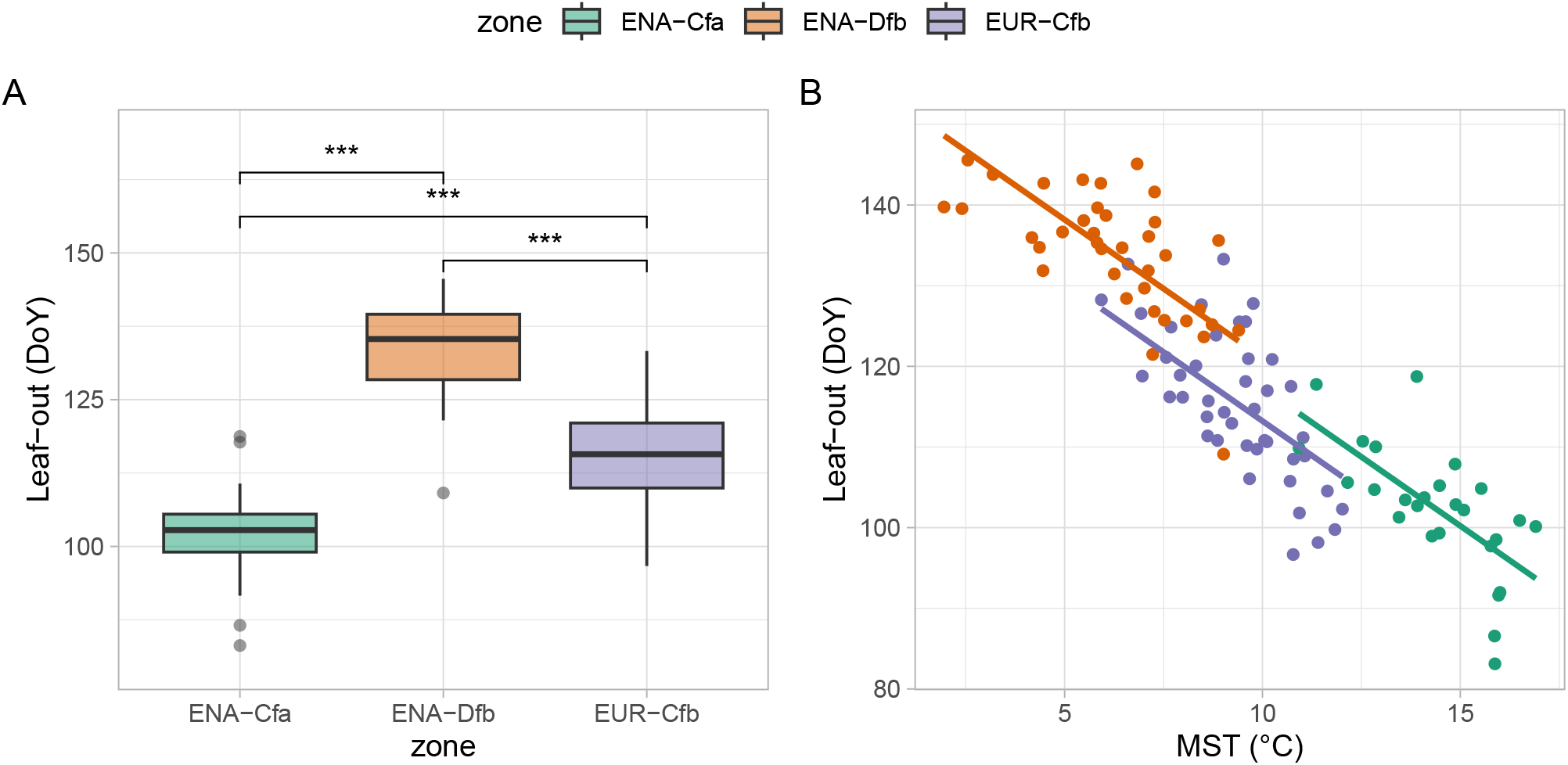
Comparing leaf-out dates in EUR and ENA. Both plots display the average leaf-out dates (day of year, DoY) determined for each site-year at the scale of the community. The data are grouped by climate zone (colors). (A) Boxplots of the site-year average leaf-out; (B) Relation of leaf-out with mean spring (March-May) temperature (MST). In (A), p-values of Wilcoxon’s tests are shown (p*<*0.001 ***, p*<*0.01 **, p*<*0.05 *, p*<*0.10 *·*, p⩾0.10 ns). In (B), linear regressions of leaf-out against MST are shown.

**Fig. 4:**
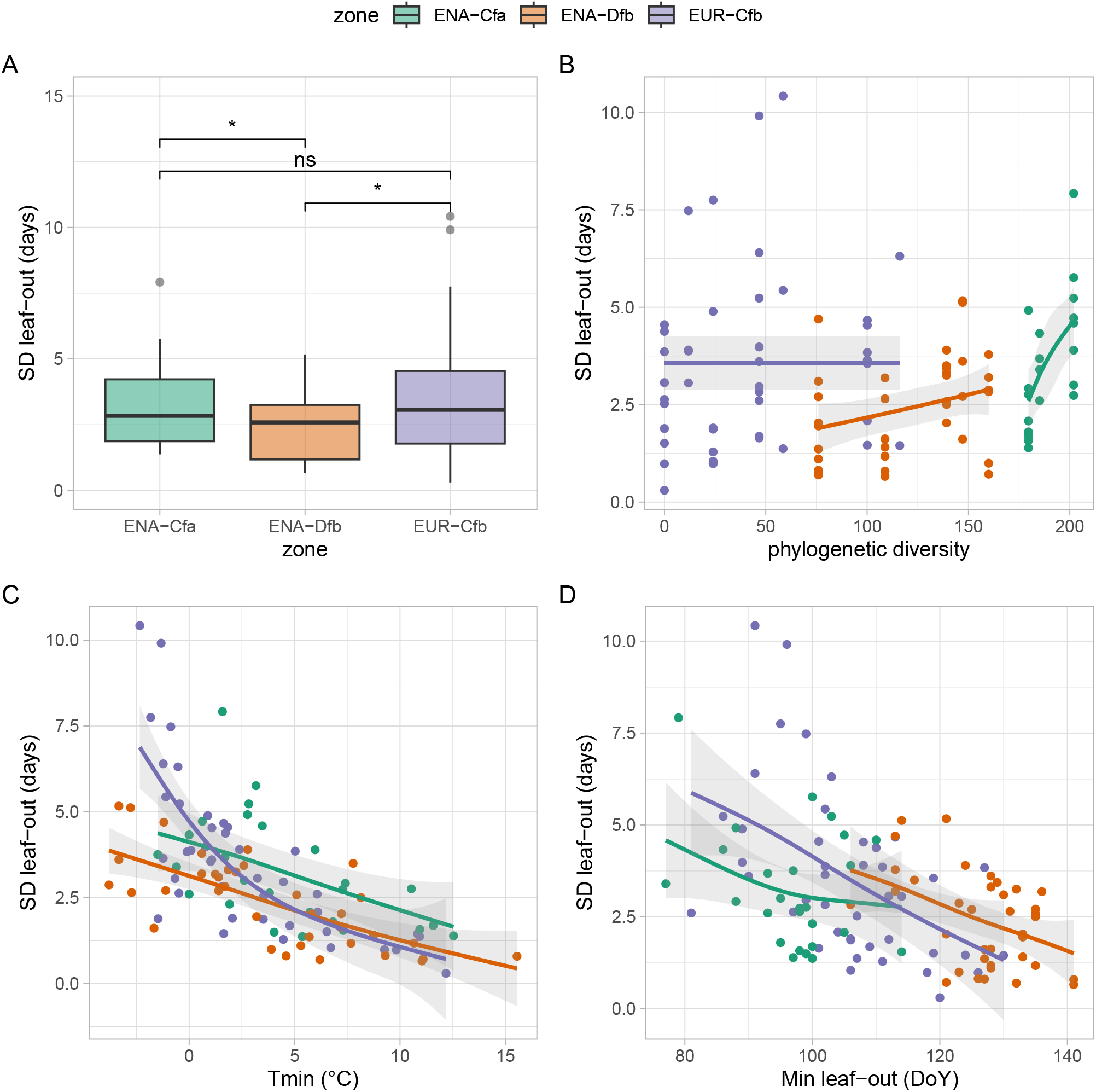
The intra-community variability of leaf-out dates in EUR and ENA. All plots display the standard deviation (SD, in days) of leaf-out dates determined for each site-year at the scale of the community. (A) Distributions of the SDs of leaf-out in EUR and ENA; Relation of SD with: (B) the phylogenetic diversity of communities, quantified as mean pairwise distance among species (see text); (C) the minimum temperature recorded over percentiles 5 to 95 of the leaf-out period (D) the minimum date of leaf-out observed on the considered site-year. In (A), p-values of Wilcoxon’s tests are shown (p*<*0.001 ***, p*<*0.01 **, p*<*0.05 *, p*<*0.10 *·*, p⩾0.10 ns). In (B), (C) and (D), the lines display fits of generalized additive models (GAMs), used here for visualization purposes. In (B), data from Bull shoals were omitted (see text).

**Fig. 5:**
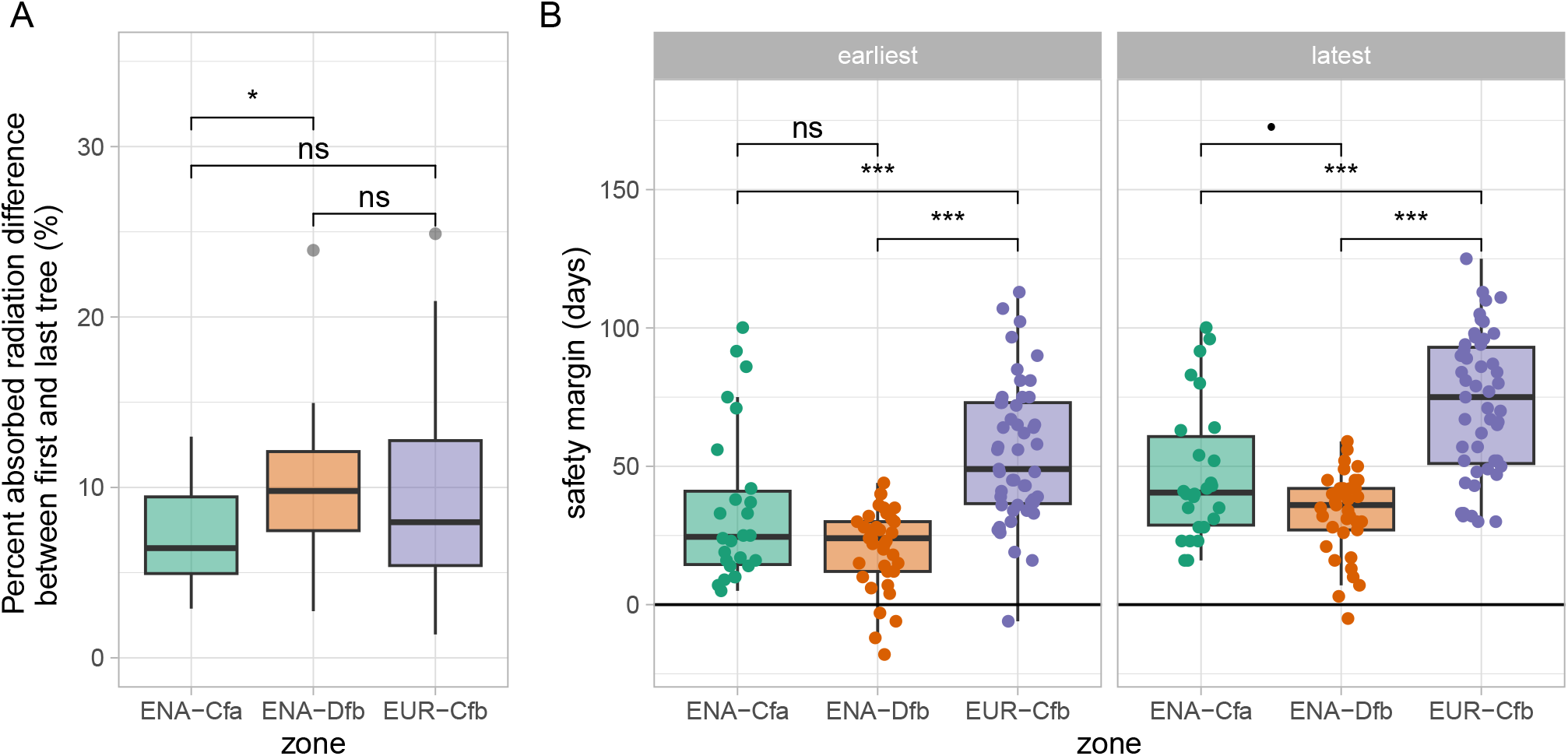
Functional impacts of the intra-community variability of leaf-out. (A) shows the relative amount of excess radiation absorbed by the earliest compared to the latest sub-ROI in the community; (B) shows the safety margin (in days) of the earliest and latest sub-ROI (two panels), as regards exposure of emerging leaves to temperatures lower than -3°C. In panel (B) data points are displayed over their respective boxplots in order to highlight the spread, and identify clearly cases with a negative safety margin.

### 2.6 Statistics

#### Comparing distributions among climate zones

We used boxplots for graphical representations of the data. They display the median, first and third quartiles (box), and go up to the largest / lowest value, no further than 1.5 times the inter-quartile range (whiskers). Data beyond the end of the whiskers are plotted individually. When comparing distributions of data among climate zones, we applied non-parametric rank sum Wilcoxon tests.

#### Statistical models

In order to answer Question 1 (see Objectives), we tested the link between the average leaf-out date and spring temperature, taking into account a possible interaction of the climate zone (i.e. relation between average leaf-out date and temperature could differ among climate zones) :

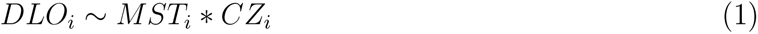

where *DLO_i_*is the average date of leaf-out (DoY) for site-year *i*, *MST* is the mean spring (March-May) temperature (°C) and *CZ* is the site climate zone.

Questions 2 and 3 were related to the influence of the community diversity and the environment on ICVLo. We first hypothesized that ICVLo, quantified as the SD of leaf-out dates, would be related to temperature and the date of leaf-out, perhaps differently across climate zones. Hence we formulated a first model:

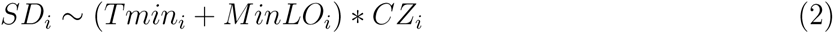

In order to test the hypothesis that more diverse communities display a higher SD, we further formulated two models:

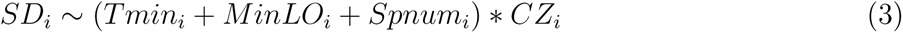

and

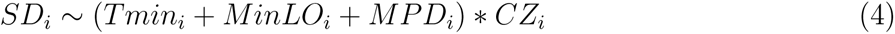

where *SD_i_*is standard deviation of leaf-out (in days) for site-year *i*, *Tmin* is the minimum temperature measured during the period extending from the 5th to the 95th percentile of the sub-ROI distribution of leaf-out (°C), *MinLO* is the date of leaf-out of the earliest sub-ROI, *Spnum* is the number of tree species contributing to the GCC signal on the phenocam ROIs, and *MPD* is a measure of the phylogenetic distance computed over those species (see above).

We used generalized linear models (GLMs) with a gamma error distribution for fitting eq. 2, 3 and 4, because the SD data were non-gaussian but followed a gamma distribution. The coefficients of a GLM relate to the mean by way of the assumed link function, which is the inverse function by default for a gamma GLM. Equations 2 to 4 include a number of terms, some of which may not be significant and hinder a proper estimation of the remaining, significant effects. In order to identify model structures including only significant terms, we used the *stepAIC* procedure in R. We compared the fitted models using the Akaike Information Criterion (AIC). Data from the Bull shoals site could not be considered for the model fitting, because we missed informations on the exact species number and identity, and therefore could not estimate the *Spnum* and *MPD* variables there.

#### Software version

All analyses were conducted with the R software (version 4.2.2) (R Core Team, 2023). Figures were plotted with the *ggplot2* package (Wickham, 2016).

## 3 Results and Discussion

### 3.1 Spring temperature across climate zones

Temperatures during the first half of the year (Jan-Jun) differed markedly among the study climate zones (Fig.1 inset). Sites in the Dfb zone (snowy climate, with warm summer) of ENA experienced negative temperatures, averaging -7°C from DoY 1 to 50, followed by a steep increase with positive temperature reached from DoY 75, and +5°C reached on DoY 100. Temperatures remained positive from DoY 1 in the Cfa zone (warm temperate climate, with hot summer) of ENA and the Cfb zone (warm temperate climate, with warm summer) of EUR, with a steeper increase from DoY 50 in Cfa, as compared to Cfb. Temperatures reached +8°C and +15°C on DoY 100 in EUR-Cfb and ENA-Cfa, respectively.

### 3.2 Comparing the leaf-out dates in ENA and EUR, and their relationships to temperature

Leaf-out dates averaged over the community spanned a larger range in ENA (from DoY 83, March 24, to DoY 146, May 26) than in EUR (DoY 97, April 7, to DoY 133, May 13) (Fig. 3). The distributions of leaf-out dates were significantly different among climate zones (pairwise Wilcoxon tests all returned p*<*0.001, Fig. 3A). Communities in ENA-Cfa leafed out on average on DoY 102 (April 12), 13 days earlier than EUR-Cfb (DoY 115, Apr 25) and 31 days earlier than ENA-Dfb (DoY 133, May 13). The three climate zones displayed spring (March-May average) temperature ranges of about 6 degrees (Fig. 3B), with the warmest spring temperatures in ENA-Cfa, and the lowest in ENA-Dfb (see also Fig. 1 inset). Noticeably, the slopes of leaf-out to spring temperature were not significantly different among zones (p*<*0.14, i.e. no interaction term was retained in eq. 1) but the intercept were, albeit marginally (Fig. 3B and Table 2: intercept of the relation was lower in EUR).

**Table 2:** Output of a linear model testing the influence of spring temperature and climate zone on the average date of leaf-out. *MST* is mean spring (March-May) temperature, *zone* identifies the climate zones.

| Parameter | Estimate | SE | t-value | p-value |
| --- | --- | --- | --- | --- |
| Intercept | 151.54 | 5.57 | 27.2 | $< 10^{-15}$ |
| MST | -3.42 | 0.38 | -9.1 | $< 10^{-14}$ |
| zone:ENA-Dfb<sup>a</sup> | 3.75 | 3.47 | 1.1 | 0.28 |
| zone:EUR-Cfb<sup>a</sup> | -4.11 | 2.46 | -1.7 | 0.10 |

<sup>a</sup> zone:ENA-Cfa set to zero

Intercontinental analyses of leaf-out dates have mostly been conducted through remote sensing studies (so-called ”start of spring” indexes) (Piao et al., 2019; Jeong et al., 2011). They evidenced trends to earlier leaf-out with recent climate change, but to our knowledge did not compare the slopes (in days per degree) that quantify the sensitivity of leaf-out to air temperature. We show here that the slopes are virtually identical in ENA and EUR, despite peculiarities of the species compositions of the European and North American floras. The rate of change of leaf-out date per unit temperature change (-3.4 days per °C, Table 2), established here across continents, is in the range of those reported for European tree species during the ”pre-season” period (from -2.0 to -4.5 days per °C, Fu et al., 2015; Zohner et al., 2018). Other works have addressed the leaf-out of plants in botanical common gardens, and showed that ENA species tend to leaf out later than EUR species when placed in the same climate conditions (e.g. Zohner et al., 2017, their Fig. 3). Results from our regression analysis are in line with this. Indeed, for the same spring temperature, EUR forests leaf out earlier than ENA forests (see Fig. 3B, and the fact that intercept of the EUR-Cfb is lower than for the ENA site, albeit marginally p*<*0.097, Table 2).

### 3.3 Comparing the intra-community variability of leaf-out dates in ENA and EUR, its dependence on species richness and environental controls

The intra-community variability of leaf-out dates (ICVLo) was in the range of 0.3 to 10.4 days, and averaged 3.1 days (3.2 days in ENA-Cfa, 2.4 days in ENA-Dfb and 3.6 days in EUR-Cfb, Fig. 4A). The mean ranks between samples of SD were significantly different among climate zones (Kruskal-Wallis test *χ*^2^=6.69, p*<*0.04), with ranks of the SD distribution being significantly lower in ENA-Dfb than in the two other zones (Fig.4A). We hypothesized that ICVLo would be larger than the intra-population variability (Fig. S1). However, the average ICVLo of 3.1 days calculated over our data (all zones grouped) was significantly lower (Wilcoxon rank sum test, p*<*0.003) than the average standard deviation of leaf-out in tree populations, that reached 4.0 days (data from Deńechère et al., 2021 consisting of 37 site-years of ground observations across 12 European tree populations). More precisely, the distribution of SD established at the community scale from phenocams in EUR was not different from the distribution of SD established at the population scale from the analysis of ground observations (Wilcoxon rank sum test, p*<*0.18, Fig. S4). How-ever, the latter was significantly higher than the distribution of SD established from phenocams in ENA (Wilcoxon rank sum test, p*<*0.001 for the comparison with ENA-Dfb and p*<*0.06 for the comparison with ENA-Cfa, Fig. S4).

Several studies have evaluated the relationship between leaf-out dates obtained from phenocams and those obtained from ground observation. These studies show that the leaf-out dates obtained from phenocams have an inter-annual amplitude very close to those obtained from ground observation (Delpierre et al., 2020; Keenan et al., 2014; Soudani et al., 2021). Hence we hypothesize that the lower SD we observed at the scale of communities, as compared to SD observed at the scale of populations (Deńechère et al., 2021), arose from an under-sampling of the actual variability in our phenocam analysis. Since we observed no systematic under-estimation of SD due to our grid-definition of sub-ROIs (Suppl. Notes S2), we hypothesize this under-sampling could arise from the fact that phenocams point mostly to dominant overstory trees, therefore overlooking inter-individual variations related to tree size (Gressler et al., 2015), developmental stage (Vitasse, 2013) or micro-environmental conditions (Pérot et al., 2021), that are captured when conducting phenological observations from the ground.

The phenocam scenes included on average more species in ENA than in EUR (median number of species was 5 in ENA-Cfa, 4 in ENA-Dfb and 2 in EUR-Cfb, Fig. S5A). This difference in species richness was also reflected in phylogenetic diversity, which was higher in ENA than in EUR (Fig. S5B). There was no significant correlation between ICVLo and phylogenetic diversity, when considering all data together (Fig. 4B, see Fig. S3 for correlation analysis). However, ICVLo tended to increase with phylogenetic diversity in ENA-Dfb (rank correlation 0.38, p*<*0.03) and ENA-Cfa (rank correlation 0.67, p*<*0.002) (Fig. 4B). In both EUR and ENA, SD decreased with warmer minimum temperatures during leaf-out (Fig. 4C), and with the date of earliest leaf-out in the community (Fig. 4D).

The model based on eq. 4, incorporating the influences of temperatures, date of leaf-out and a measure of the phylogenetic diversity of the community fitted the ICVLo data best (Table 3). The terms of the full model were generally coherent with the visual depiction from Fig. 4, considering the model was fitted with an inverse link function (see Methods). For instance, the coefficient estimates for variables Tmin and MinLO were positive in the model (Table 4), coherent with the negative correlation of SD with those variables (Fig. 4C,D). The interaction term Tmin*EUR-Cfb was positive, indicating that the response of SD to temperature in EUR-Cfb was more pronounced (i.e. with a steeper negative slope) than in ENA-Cfa and ENA-Dfb (see Fig. 4C). The interaction terms MPD*EUR-Cfb and MPD*ENA-Dfb were both positive, indicating that the response to MPD was weaker there than in ENA-Cfa (see Fig. 4B).

**Table 3:**
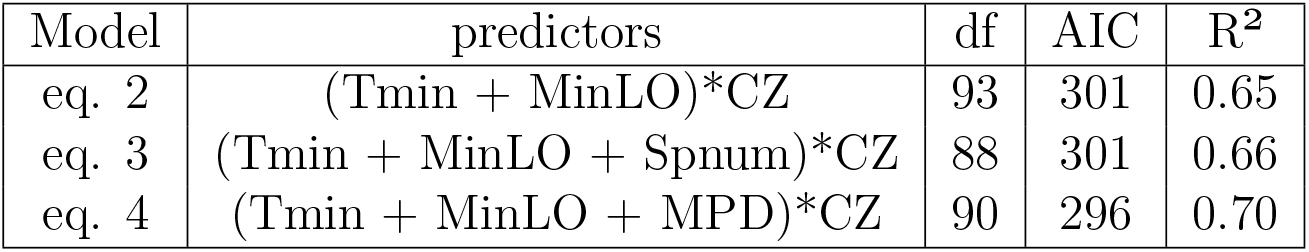
Comparing models fitted to the ICVLo data. *df* is the number of degrees of freedom, obtained after model structure simplification by the *stepAIC* procedure. *AIC* is the Akaike Information Criterion. See text for the description of models.

**Table 4:**
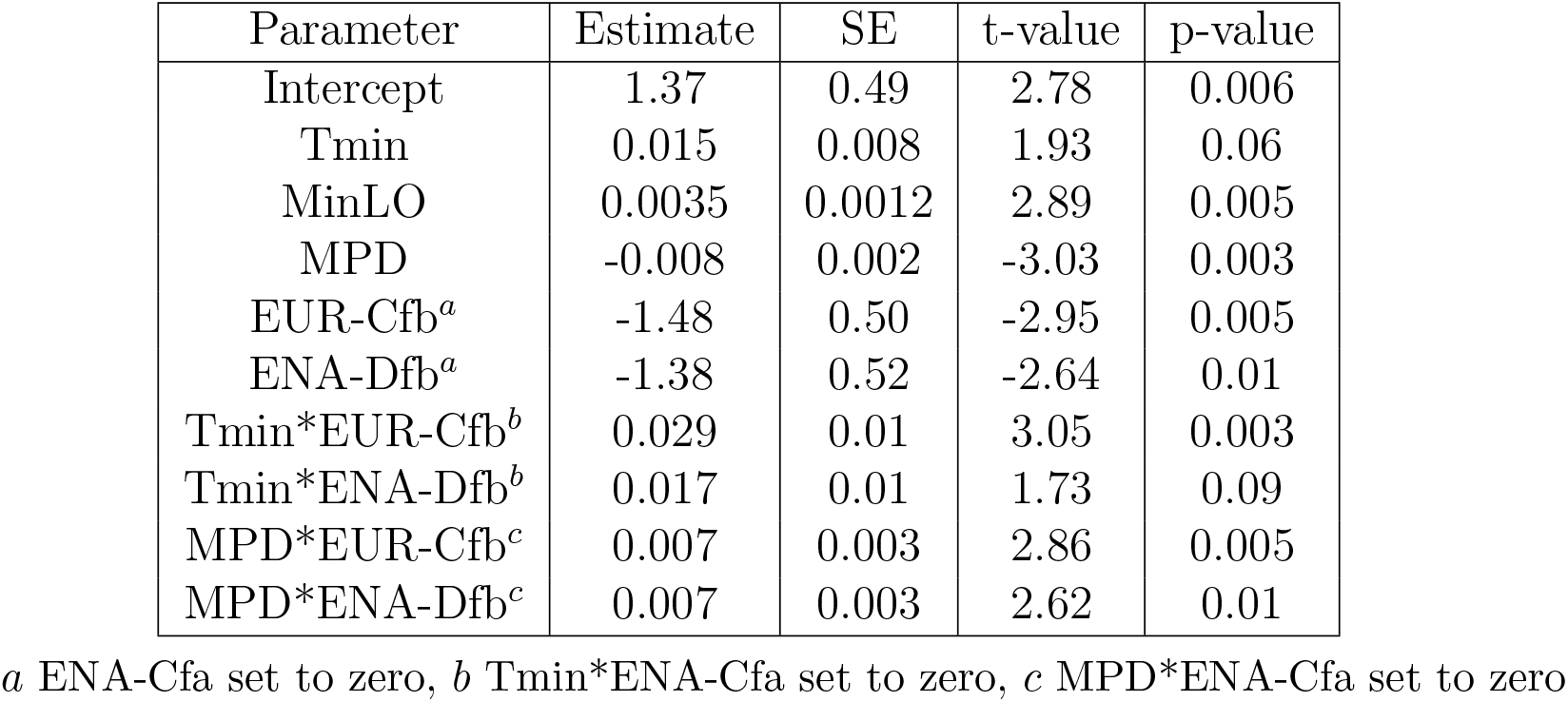
Summary of the best Generalized Linear Model fit to the SD data. The model (eq. 4) was fitted with a gamma error distribution, using an inverse link function. Only terms of the models retained by the *stepAIC* procedure are displayed here. *SE* is the standard error of the parameter, *MinLO* is the minimum leaf-out date, *Spnum* is the species number, *MPD* is a measure of the phylogenetic diversity in the community. See text for details.

The correlation of ICVLo with minimum temperatures occurring during the leaf-out period (Fig. 4C) mirrors earlier results obtained at the scale of tree populations (Deńechère et al., 2021). Warmer springs have been shown to hasten the speed of leaf-out across scales, from the scale of the bud (Basler and Körner, 2014), to the individual tree crown (Deńechère et al., 2021), to the population (Deńechère et al., 2021). Here, we show that this result extends to the scale of the community. Recently, Lin et al. (2023) developed a model simulating the progress of leaf-out at the scale of a tree population. In this model, each individual tree has a particular sum of forcing temperatures to reach for leaf-out to occur. In line with our results, the model predicts a longer period of time from the first to last tree to leaf-out (hence extended intra-specific variability) during cooler springs (see also Fig. S1 of Deńechère et al., 2021).

A study conducted at the scale of Germany showed that warmer springs resulted in a ”loss of phenological synchrony” (i.e. higher variability of leaf-out) among populations of European Beech trees (Zohner et al., 2018). Our results could appear to contradict this work, because we found a negative link between ICVLo and temperature during leaf-out (i.e. warmer temperatures during leaf-out are associated with smaller ICVLo, that is a higher phenological synchrony in the communities). This contradiction disappears once we take into account the fact that these two studies do not consider the effect of temperatures in the same time window. We related ICVLo to the temperature conditions occurring during the period of leaf-out (Fig. 4C) while Zohner et al. 2018 considered temperature conditions occurring during the 60 days preceding the average date of leaf-out. Similar to Zohner et al. 2018 (their Fig. 1E), we found that early leaf-out (caused by a high ”pre-season” temperature) was associated with a higher variability of leaf-out (Fig.4D). However, the influence of the date of leaf-out on ICVLo was of second order, as compared to the influence of temperature conditions during leaf-out (i.e. partial correlations of SD with temperature, controlling for leaf-out date, were stronger than partial correlations of SD with date, controlling for temperature, see Table 5).

**Table 5:**
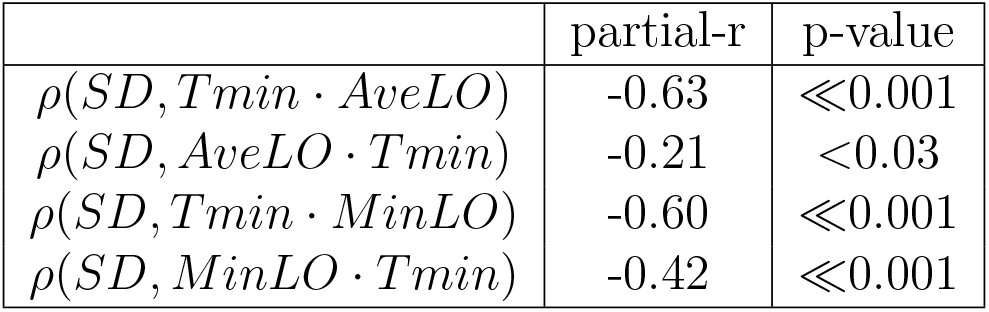
Partial correlation of SD with temperature and leaf-out date. *AveLO* is the average leaf-out date, *MinLO* is the minimum leaf-out date. For example, *ρ*(*SD, Tmin · AveLO*) is the partial correlation of SD with Tmin, controlling for AveLO.

### 3.4 Ecological implications of the intra-community variability of leaf-out

Individual trees leafing-out first in the community received on average 8% more radiation (from 7% in ENA-Cfa to 10% in ENA-Dfb) over spring than the last trees leafing-out (remembering we focused here on the dominant overstory trees) (Fig. 5A). Under the naive hypothesis that photosynthesis scales with incoming radiation, as in a simple light-use efficiency (LUE) model (Baldocchi and Peñuelas, 2019), and that NPP (net primary productivity, or biomass productivity) is proportional to GPP (gross primary productivity, the gross amount of carbon fixed by photosynthesis) (Collalti and Prentice, 2019), and ignoring intraspecific and leaf age variation in LUE, this would straightforwardly translate in a 8% difference in tree growth. This potential 8% difference is much smaller than the variability observed in basal area increment of tree growth we compute, e.g. at the Barbeau (FR-Fon) forest (Table 1) for which we could access individual tree growth data (coefficient of variation of tree basal area increment, normalized by crown area was 35% among dominant trees). Along with other evidence (e.g. the fact that phenological rank in tree populations are not systematically correlated with growth, Delpierre et al., 2017), this points to a second-order influence of leaf phenology in determining the inter-individual growth of trees (see also Cufar et al., 2015 and de Sauvage et al., 2022).

The safety margin against exposure to frost was significantly smaller in ENA than in EUR (Fig. 5B). As expected considering the trend to increasing temperatures in spring, early sub-ROIs had a smaller safety margin than late sub-ROIs, whatever the climate zone (Fig. 5B, average safety margins in the ”earliest” sub-ROI were smaller than in the ”latest” sub-ROI). In EUR, we detected 1 site-year (i.e. 2% of the EUR data, Table 6) for which the earliest sub-ROI was exposed to a frost below -3°C. In ENA, this rose to 4 site-years (6% of the ENA data), belonging to two sites, both located in the cooler ENA-Dfb zone (Fig. 1 inset). This result is consistent with calculations of a higher probability of frost exposure for North American, as compared to European species (Zohner et al., 2020), and suggests that North American temperate forests, indeed, experience late frosts more frequently than European forests. Not only the frequency, but also the extent of exposure to frost of ENA vs. EUR forests differed in our dataset. When late frost struck ENA forests, it could affect a large proportion of the community (Table 6).

**Table 6:** Percent of the community affected by late spring frost. We report here the percentage of sub-ROIs that had already leafed-out when late frost (Tmin*<*-3°C) occurred.

| Zone | Site | Year | Tmin ($^{\circ}\text{C}$) | Percent frost (%) |
| --- | --- | --- | --- | --- |
| ENA-Dfb | Arbutus Lake | 2009 | -3.3 | 56 |
| ENA-Dfb | Arbutus Lake | 2010 | -3.3 | 74 |
| ENA-Dfb | Arbutus Lake | 2013 | -4.4 | 1 |
| ENA-Dfb | Sanford | 2020 | -3.8 | 90 |
| EUR-Cfb | Barbeau | 2021 | -3.0 | 1 |

## Conclusions and perspectives

We were able to characterize the intra-community variability of leaf-out dates through the analysis of images acquired automatically by phenological cameras. This methodological achievement is a promising step in the process of characterizing the current and past variability of ecological traits in tree communities (e.g. the grid-based analysis of images could be applied to other ecological traits such as detecting variations in the health status, differential attacks by herbivores etc.) with implications on the forest biocenosis (e.g. the timing of leaf emergence determines the availability of food for many herbivorous insect species). It complements other attempts to characterize the variability of phenology at the scale of a landscape, using UAVs (Klosterman et al., 2018; Berra et al., 2019) or satellite data (Moon et al., 2022).

We found ample ICVLo at all study sites. The ICVLo was comparable in ENA and EUR, despite ENA forests being more species-rich and phylogenetically diverse, pointing to a stronger environmental than biotic control of ICVLo. The ecological consequences of ICVLo are at least two-fold: (1) on average, trees that leaf-out the earliest in a community were exposed to about 8% more radiation over spring than the latest trees, (2) those early trees have a lower safety margin regarding exposure to late frost, and the safety margin was smaller in ENA as compared to EUR. Further ecological consequences of ICVLo remain to be explored, for example the impact of ICVLo on soil water content in a context of drier environmental conditions. The approaches developed in this work could also be extended to the analysis of the intra-community variability in leaf senescence, which is even greater than ICVLo (Delpierre et al., 2017) and whose determinants are probably more complex.

## Supporting information

Supplementary informations

## Acknowledgements

We thank our many collaborators, including site PIs and technicians, for their efforts in support of this study, and the PhenoCam network. In particular, we thank PIs from the European Integrated Carbon Observation System (ICOS) for providing the meteorological data and site information. We thank Sara DiBacco Childs (Duke Forest Executive Director) and Blake Tedder (Assistant Director of Engagement, Duke University) for their contribution. ND acknowledges funding from ANR (TAW-tree project, grant #ANR-23-CE01-0008-01).The development of Phe-noCam has been funded by the Northeastern States Research Cooperative, NSF’s Macrosystems Biology program (awards EF-1065029 and EF-1702697), and DOE’s Regional and Global Climate Modeling program (award DE-SC0016011). JL was funded by the China Scholarship Council (grant #202008330320). AK and FK were funded by the German Federal Ministry of Education and Research (BMBF) as part of ICOS, by the Deutsche Forschungsgemeinschaft (INST 186/1118-1 FUGG) and by the Ministry of Lower-Saxony for Science and Culture (DigitalForst: Niedersächsisches Vorab (ZN 3679)). MW acknowledges funding from the UK Forestry commission. AN acknowledges funding from DOE Ameriflux Network Management Project, DOE NICCR 08-SC-NICCR-1072, DOE-TES 11-DE-SC-0006700, USDA Forest Service Eastern Forest Environmental Threat Assessment Center 08-JV-11330147-038 and 13-JV-11330110-081. JCD acknowledges funding from DOE DE-SC0023309, PHydrauCC #ANR-21-CE02-0033-02. JWM acknowledges funding from DOE DE-AC02-05CH11231, NSF LTER DEB-1832210. SOD acknowledges funding from NIFA (award #2021-67034-35129). KD acknowledges funding from National Science Foundation DEB-2044818 and USDA Hatch Project #MICL02716. KMH and GC were funded by the JABBS Foundation and the University of Birmingham. ARD acknowledges funding from DOE Ameriflux Network Management Project award to ChEAS core site cluster.

