## Supplementary informations for "Phenology across scales: an intercontinental analysis of leaf-out dates in temperate deciduous tree communities"

Nicolas Delpierre et al.

*Supplementary material*

**Supplementary Notes S1. The amplitude of leaf-out increases as one moves from the scale of the population to the scale of the community**.

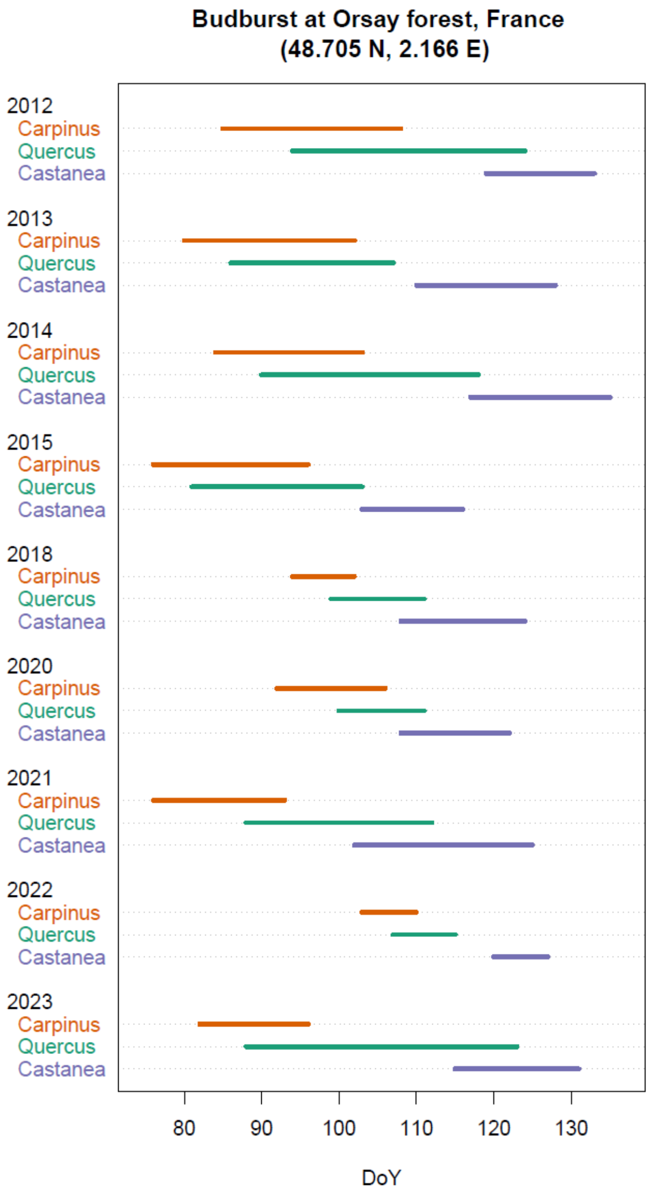


**Figure S1. Variability of leaf-out in a forest community.** For each species and year (2012 to 2023), the lines connect the minimum and maximum date of leaf-out observed across 60 individuals (per species). All data were acquired at the Orsay forest, France (48.705 N, 2.166 E). These three species are the main of a community of about 10 deciduous tree species in the Orsay forest. *Carpinus betulus*, L. is an understory species. *Quercus petraea*, (Matt.) Liebl. and *Castanea sativa*, Mill. are overstory species.

**Table S1. Species number and names at each site.** We report here the number ("Species number") and name of deciduous species identified in the site-specific phenocam ROIs delineated for this study.

| Zone | Site | Species number | Species identity and contribution to the canopy |
|---|---|---|---|
| EUR | Alice Holt | 2 | *Quercus robur* (70%), *Fraxinus excelsior (30%)* [share of canopy area, estimated visually] |
| EUR | Barbeau (FR-Fon) | 2 | *Quercus petraea* (75%), *Carpinus betulus (25%)* [share of canopy area, estimated visually] |
| EUR | DE-Hai (DE-Hai) | 3 | *Fagus sylvatica (66%), Fraxinus excelsior (26%), Acer pseudoplatanus (8%)* [share of basal area, from Mund et al. 2010] |
| EUR | Hohes Holz (DE-HoH) | 3 | *Quercus petraea (48%), Fagus sylvatica (39%), Carpinus betulus (13%)* [share of basal area, from Holtmann et al. 2021] |
| EUR | Leineflede (DE-Lnf) | 1 | *Fagus sylvatica* (100%) [share of canopy area, estimated visually] |
| EUR | Soroe (DK-Sor) | 1 | *Fagus sylvatica* (100%) [share of canopy area, estimated visually] |
| EUR | Hesse (FR-Hes) | 4 | *Fagus sylvatica (88%), Quercus robur (3%), Quercus petraea (1%), Carpinus betulus (8%)* [share of basal area, pers. comm. from M. Cuntz] |
| EUR | Mill Haft | 2 | *Quercus robur (95%), Betula pendula (5%)* [share of canopy area, estimated visually] |
| ENA | Alligator River (US-NC4) | 2 | *Nyssa aquatica (85%), Taxodium distichum (15%)* [share of canopy area, estimated visually] |
| ENA | Arbutus lake | 5 | *Acer saccharum (31%), Fagus grandifolia (15%), Acer rubrum (10%), Betula alleghaniensis (35%), Prunus serotina (9)* [share of canopy area, estimated by individual crown delineations on phenocam images] |
| ENA | Bull shoals | > 9 | *Quercus alba, Quercus stellata, Quercus velutina, Quercus falcata, Carya cordiformis, Carya tomentosa, Fraxinus americana, Diospyros virginiana, Cercis canadensis* [exact species composition and percent contribution to ROI could not be determined] |
| ENA | Downer woods | 4 | *Fraxinus americana (45%), Tilia americana (45%), Quercus alba (5%), Quercus rubra (5%)* [share of canopy area, estimated visually] |
| ENA | Duke hw (US-Dk2) | 5 | *Liquidambar styraciflua (46%), Liriodendron tulipifera (15%), Carya tomentosa (29%), Quercus alba (5%), Quercus michauxii (5%)* [share of canopy area, estimated by individual crown delineations on phenocam images] |
| ENA | Harvard Forest (US-Ha1) | 4 | *Quercus rubra (75%), Acer rubrum (15%), Betula alleghaniensis (5%), Fagus grandifolia (5%)* [share of canopy area, estimated visually] |
| ENA | Morgan monroe (US-MMS) | 5 | *Liriodendron tulipifera (56%), Acer saccharum (19%), Ulmus americana (6%), Juglans nigra (6%), Sassafras albidum (13%)* [share of number of trees, estimated by individual tree identification on phenocam images] |
| ENA | Sanford | 6 | *Acer saccharum (30%), Tilia americana (26%), Ostrya virginiana (4%), Fagus grandifolia (26%), Carya cordiformis (9%), Quercus rubra (4%)* [share of number of trees, estimated by individual tree identification on phenocam images] |
| ENA | Willow creek (US-WCr) | 4 | *Acer saccharum (68%), Tilia americana (5%), Fraxinus pennsylvanica (26%), Quercus rubra (1%)* [share of number of trees, calculated from field inventory] |

**Supplementary Notes S2. Quantifying the uncertainty on the estimation of the standard deviation of leaf-out at the scale of the community from a grid-based approach**.

We aimed to quantify the uncertainty on the estimation of the standard deviation of leaf-out at the scale of the community (SD) caused by the use of a grid-based definition of sub-ROIs. For this, we compared the value of SD determined in one site-year under two cases for the identification of sub-ROIs: (1) sub-ROIs corresponding to the exact delineation of individual tree crowns (that will be our reference) and (2) sub-ROIs determined automatically by the application of the grid approach (Fig. S2.1).

The comparison of SD determined in the two cases is shown in Fig. S2.2. The root mean squared difference of SD is 0.36 days.

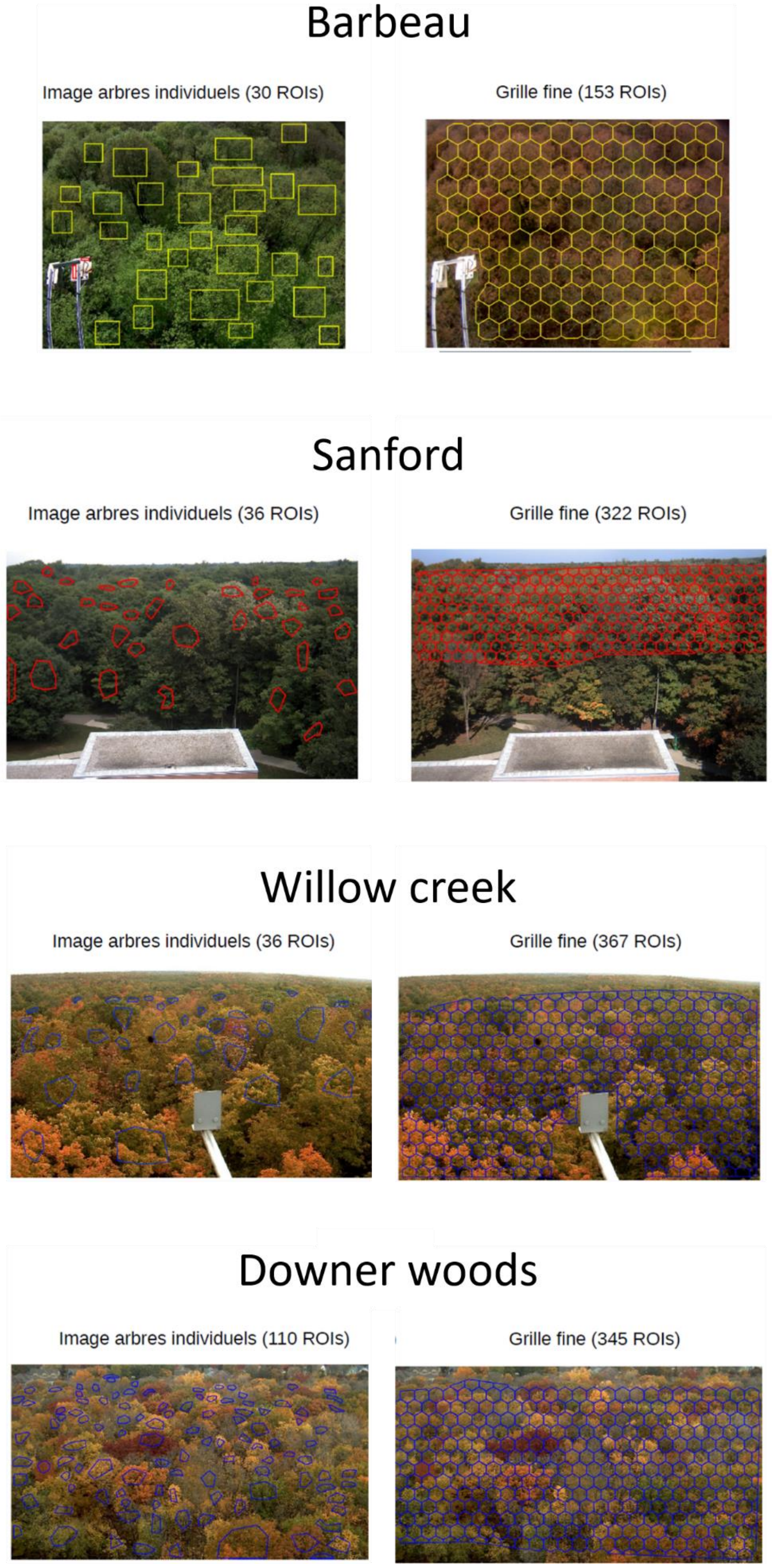


**Figure S2.1**. Phenocam scenes displaying the individual ROIs used to determine the intra-community variability of leaf-out in two approaches: the individual tree (left column) and the systematic grid (right column).

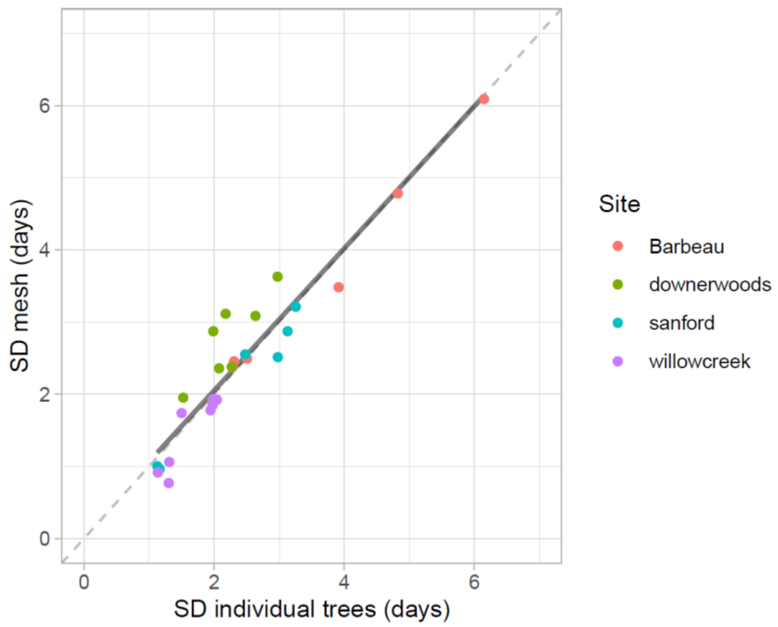


**Figure S2.2**. Standard deviation (SD) of leaf-out determined by the systematic grid approach as a function of SD determined using the individual tree approach. Each data point represents a site-year. The overall fit to the data appears as a solid line. The identity line is dashed.

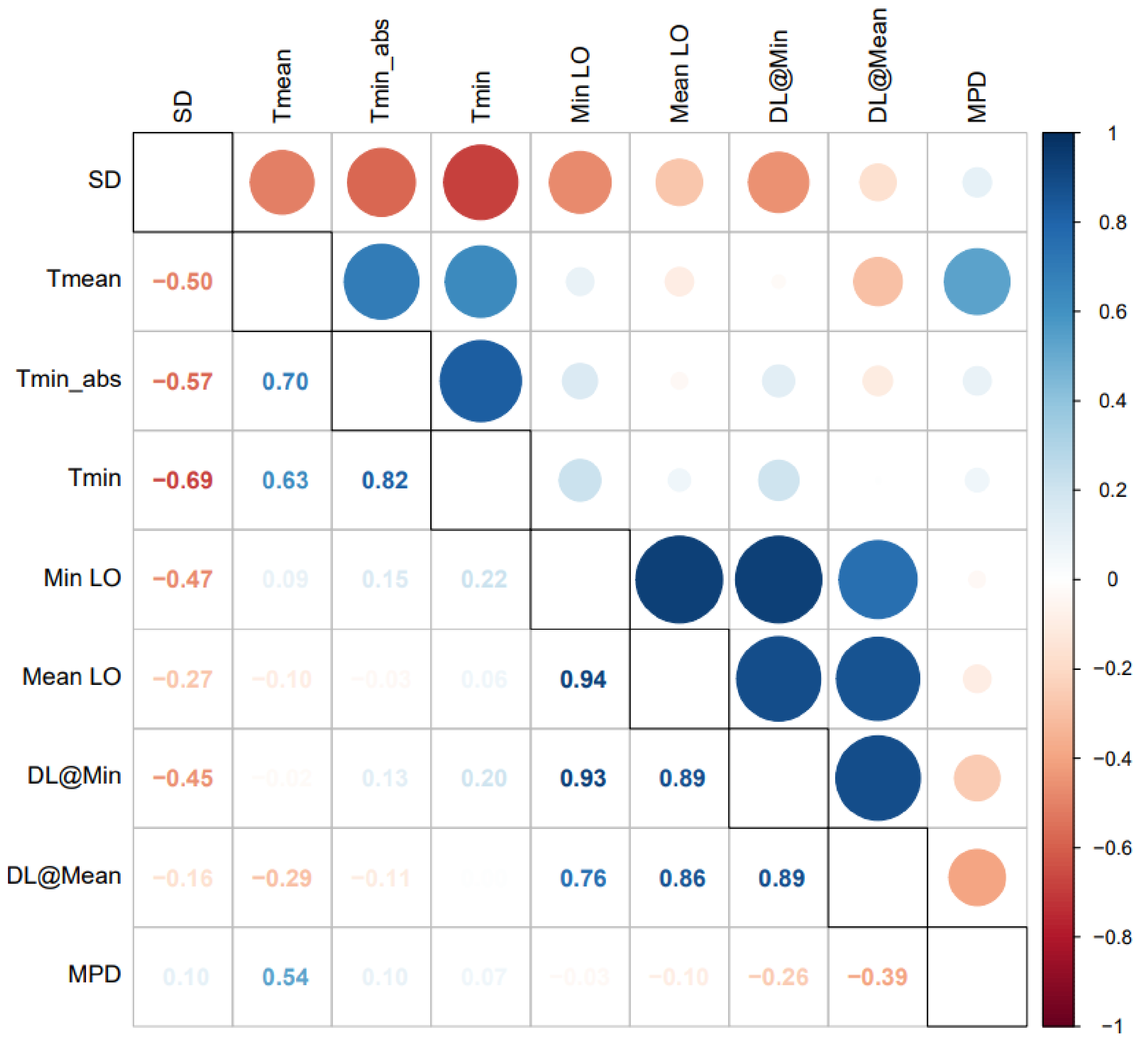


**Figure S3. Correlation matrix evidencing the dependence of the intra-community of leaf-out (SD) to phenological, environmental and phylogenetic variables.** *Tmean* = average temperature during the leaf-out period (defined as the time interval from the first to the last sub-ROI to leaf-out), *Tmin*_abs = absolute minimum temperature during the leaf-out period, *Tmin* = minimum temperature during the period extending from the 5th to the 95th centile of the sub-ROI distribution of leaf-out, *Min LO* = date of the leaf-out for the earliest sub-ROI, *Mean LO*= average date of leaf-out across sub-ROIs, *DL@Min* = daylength at the *Min LO* date, *DL@Mean* = daylength at the *Mean LO* date, *MPD* = mean pairwise distance representing the phylogenetic variability in the community. The matrix displays results of Spearman's rank correlation tests.

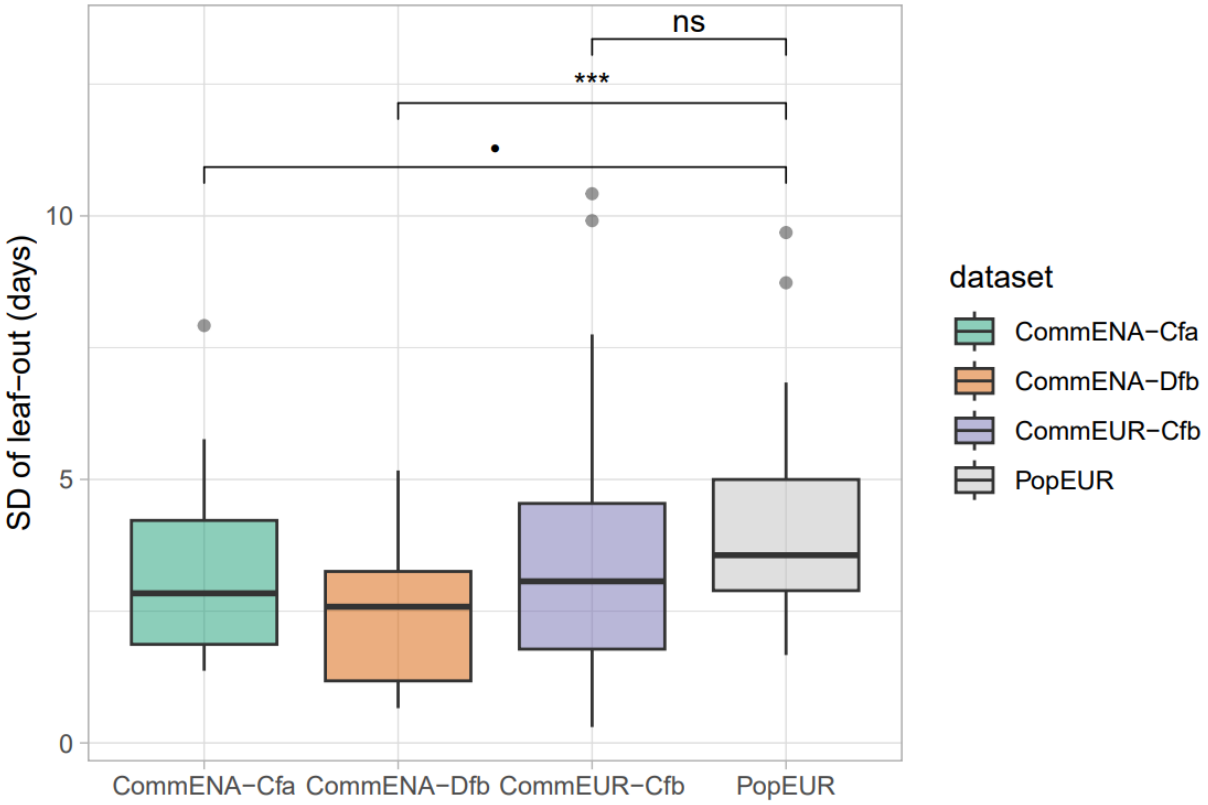


**Figure S4. Distributions of the standard deviation of leaf-out observed at the scale of the population and of the community.** The plot compares the distributions of the standard deviation of leaf-out obtained at the scale of the community through phenocam (colored boxplot; the ones analysed in the main text, see e.g. Fig. 4A) with data obtained at the scale of the tree population from ground observation (grey boxplot, data previously published in Denéchère et al. 2021). P-values of Wilcoxon's tests are shown ($p<0.001$ ***, $p<0.01$ **, $p<0.05$ *, $p<0.10$ •, $p\geq0.10$ ns).

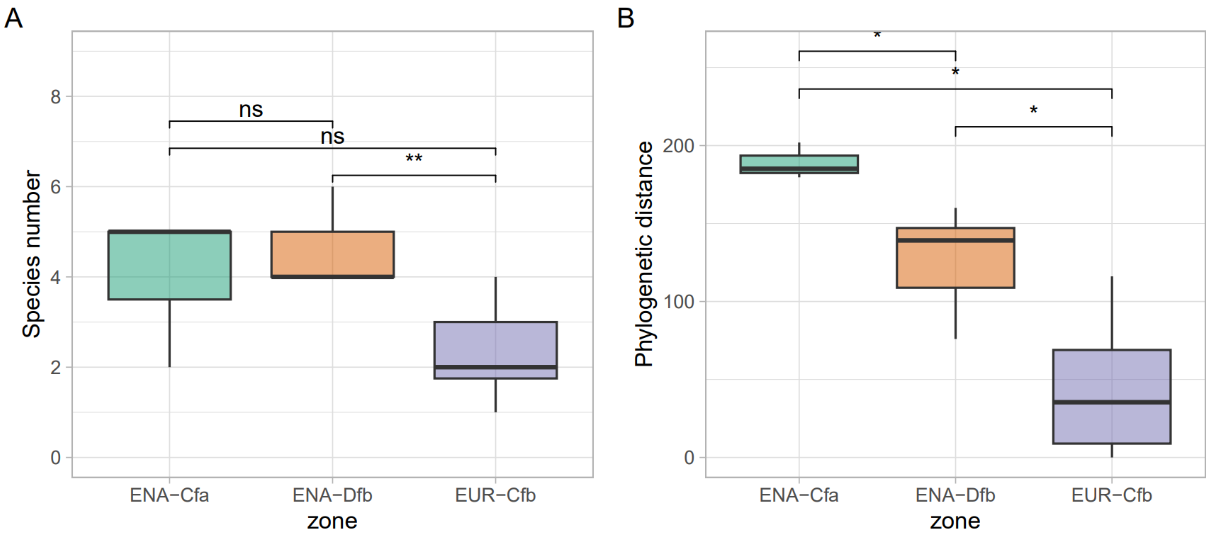


**Figure S5. Canopy species number and phylogenetic distance in EUR and ENA communities.** (A) displays the number of tree species identified in the regions of interest (ROIs, see main text) delineated from the phenocam scenes. (B) displays the *mean pairwise distance*, a measure of phylogenetic distance in the communities (see main text for details.). Both plots contain unique data for each site (i.e., representation of sites are not weighted by the number of site-years, which varies substantially among sites, see Table 1 in main text). P-values of Wilcoxon's tests are shown ($p<0.001$ ***, $p<0.01$ **, $p<0.05$ *, $p<0.10$ •, $p\geq0.10$ ns).
